# CROWN: Curated Repository Of Well-resolved Non-covalent interactions

**DOI:** 10.64898/2026.03.30.714168

**Authors:** Robin Poelmans, Wout Van Eynde, Bence Bruncsics, Balint Bruncsics, Adam Arany, Yves Moreau, Arnout Voet

## Abstract

The development of machine-learning models for protein–ligand interactions is constrained by the quality and diversity of the available structural data. Existing resources force researchers into a trade-off: carefully curated collections such as PDBBind and HiQBind offer high structural reliability but cover only a narrow slice of the Protein Data Bank (PDB), whereas large-scale resources such as PLINDER provide broad coverage with minimal quality control. We present CROWN (Curated Repository Of Well-resolved Non-covalent interactions), a machine-learning–ready dataset that reconciles scale and rigor through a fully automated preprocessing pipeline. Starting from the PDB database, CROWN applies a series of interleaved quality filters and processing stages that address crystallographic resolution, ligand identity, pocket completeness, structural repair, interaction quality, and protonation at physiological pH. The pipeline finishes with a constrained energy-minimization step built on custom flat-bottomed restraints — a step absent from all existing protein–ligand datasets — that balances crystallographic evidence against the relaxation of intramolecular strain. By reconciling the heterogeneous refinement practices of different depositions without distorting the experimentally observed binding geometry, this step yields a structurally uniform collection of 178,263 complexes, representing a roughly four-fold increase in protein diversity over PDBBind and HiQBind. Rather than organizing the data around sparsely available, bias-prone binding affinities, CROWN adopts a geometry-centric design philosophy that treats the three-dimensional arrangement of atoms at the binding interface as a self-consistent source of information. To demonstrate its value as a training resource, we trained two knowledge-based scoring functions on CROWN and benchmarked them on CASF-2016: relative to HiQBind-trained counterparts, CROWN-trained models showed markedly improved ranking power (mean Spearman correlation rising from 0.509 to 0.637) and docking power (top-1 near-native pose recovery of 0.785 versus 0.724). Because CROWN imposes no requirement for affinity labels, it can in principle support any model that learns from or is evaluated against protein–ligand complex structures. We anticipate that it will serve as a broadly useful resource for tasks such as the training of binder generation, protein design or protein folding models conditioned on bound ligands, the development of scoring functions or benchmarking of interaction-prediction methods.

## Introduction

Machine learning is reshaping structural biology. Protein folding models have redefined the accuracy ceiling for structure prediction, and equivariant diffusion models now generate realistic protein–ligand binding poses [1, 2]. The performance of such models depends critically on the quality and breadth of their training data, yet in protein–ligand structural biology the available datasets force researchers into a compromise between rigor and scale.

The rigorous end of this spectrum was defined nearly two decades ago by the Astex Diverse Set [3], which established a template for what a trustworthy protein–ligand dataset should demand. Rather than accepting deposited coordinates at face value, Hartshorn et al. reconstructed the electron density for each ligand from the deposited structure factors and retained only complexes whose binding-site atoms were unambiguously supported by the experimental evidence. Each entry was meticulously prepared: crystallization additives and other non-relevant species were removed, bond orders and connectivity were assigned explicitly, protonation and tautomeric states were resolved, and structures were assessed for steric clashes and other physical implausibilities. The resulting 85 diverse, drug-like, high-resolution complexes remain a reference point for structural fidelity, but their number reflects the cost of (semi-)manual curation. Modern curated resources inherit this philosophy while pushing on scale: HiQBind [4], for instance, subjects each complex to extensive and often manual preprocessing—modeling missing protein atoms, assigning correct bond orders, and resolving protonation states—producing structures reliable enough for tasks where geometric fidelity matters. The cost, again, is coverage: manual or semi-automated curation limits such collections to at most a few tens of thousands of entries, a modest fraction of the more than 200,000 complexes currently deposited in the PDB. As a consequence, they under-represent the chemical and biological diversity of known protein–ligand interactions, which can impair the generalization of models trained on them.

At the other end, PLINDER [5] systematically catalogs protein–ligand systems across the entire PDB, assembling nearly 650,000 entries that span a far broader range of protein families, organism sources, and ligand chemotypes than earlier resources. PLINDER, however, applies only minimal quality filtering and largely retains structures in their deposited form, leaving common artifacts uncorrected. For data-hungry methods such as deep generative models or neural network–based scoring functions, unresolved atoms, steric clashes, and incorrect bond assignments introduce systematic noise into training and undermine model reliability.

SPINDR [6] represents an important step toward reconciling quality with scale, filtering PLINDER for drug-like small-molecule ligands and applying structural preprocessing such as modeling missing protein atoms and assigning bond orders and protonation states.

In this paper, we introduce CROWN (Curated Repository Of Well-resolved Non-covalent interactions), a machine learning–ready structural dataset that bridges the gap between quality and quantity. CROWN carries the quality-control philosophy pioneered by the Astex Diverse Set and automates it across the full breadth of the PDB. Building on SPINDR, CROWN extends its scope along four axes: we broaden the coverage beyond single protein–single ligand pairs to include binding sites with organic and inorganic cofactors; we consider complexes determined by cryo-EM in addition to X-ray crystallography, widening the structural coverage of the dataset; we construct the dataset directly from the deposited mmCIF structures rather than inheriting upstream annotations, allowing each complex to be validated independently; and we release the full pipeline as open-source software so that others can reproduce and extend our work. CROWN thus retains the broad scope of PLINDER while subjecting every protein–ligand complex to a fully automated pipeline of quality filtering, structural repair, protonation, and constrained energy minimization. This final minimization step employs custom flat-bottomed restraints designed to balance fidelity to the experimental evidence against the relaxation of inter- and intramolecular strain, respecting experimental coordinate uncertainty while correcting physically implausible contacts [7, 8]. To our knowledge, this systematic application of constrained energy minimization is unique among currently available protein–ligand interaction datasets, and it is what allows CROWN to reconcile scale with structural uniformity.

Like PLINDER and SPINDR, CROWN moves away from the affinity-label paradigm that organizes established benchmarks such as PDBBind [9]. Affinity data exist for only a subset of structurally characterized complexes, so an affinity-centric framing necessarily excludes many informative structures. CROWN instead adopts a geometry-centric design: rather than organizing the dataset around measured binding constants, it treats the three-dimensional arrangement of atoms at the protein–ligand interface—the interaction geometry itself—as a rich, self-consistent source of structural information that requires no external labels.

We envision CROWN as a broadly useful resource for the structural biology, cheminformatics, and machine-learning communities. Its combination of scale, structural quality, and geometry-centric design makes it well suited for tasks such as training models for binder generation [2], ligand-conditioned protein design [10], or complex structure prediction [11], as well as developing knowledge-based scoring functions and statistical potentials [12]. In what follows, we describe the design and benchmarking of CROWN, characterize its scope relative to existing resources, and release the full dataset through a freely accessible web interface.

## Methods

The CROWN pipeline applies five sequential curation and filtering steps, described below in pipeline order and summarized in Figure 1.

**Figure 1:**
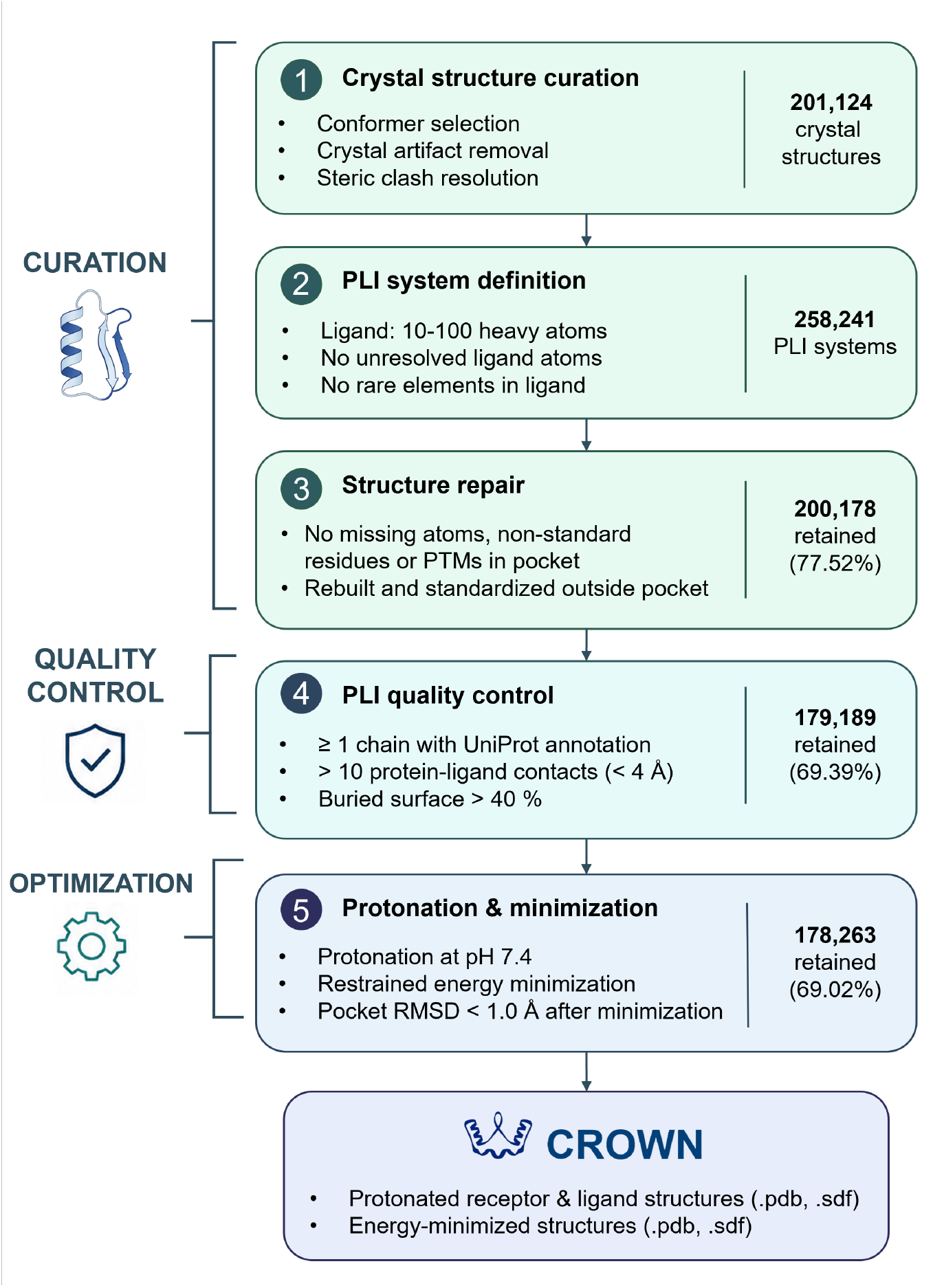
The CROWN preprocessing pipeline. Starting from 201,124 experimental structures, the pipeline applies curation (green), quality control (teal) and optimization steps (blue) in the order shown, forming a staged attrition funnel. For quality control both data (RSR:real-space R-value, RSCC: real-space correlation coefficient) and interaction quality were assessed. The number of systems passing each stage and the corresponding pass rate are indicated on the right. The final CROWN dataset comprises 178,263 complexes, distributed as protonated and energy-minimized receptor (.pdb) and ligand (.sdf) files.

### Step 1: Crystal structure curation

The CROWN preprocessing pipeline operates on mmCIF files from the Protein Data Bank (PDB) for all X-ray crystallography and cryo-electron microscopy structures with a resolution better than 3 Å. For each structure, alternative conformers were collapsed to the maximum-occupancy state using Gemmi [13] and common artifacts (as enumerated in PLINDER [5]) were removed. Inter-chain steric clashes — defined as heavy-atom contacts below 1.8 Å — were resolved based on severity: isolated single-atom clashes were interpreted as missing covalent bonds, whereupon the two clashing chains were merged accordingly, while more extensive overlaps were resolved by discarding the smaller of the two chains.

### Step 2: Protein-Ligand interaction system definition

Within each cleaned structure, candidate ligand chains were identified as chains containing between 10 and 100 heavy atoms and at least two carbon atoms. Because covalent ligands were merged with their protein chains in Step 1, they are automatically removed by this size criterion. Each candidate was validated against the Chemical Component Dictionary (CCD) residue definitions to flag unresolved heavy atoms [14]; only terminal oxygens lost to inter-residue bonds were tolerated. Chains passing this check defined the ligand of a protein–ligand interaction (PLI) system. The surrounding environment was then assembled by selecting all chains within a 3 × 3 × 3 crystallographic unit-cell box that contained at least one heavy atom within 4 Å of any ligand heavy atom present in the central unit-cell box, ensuring that symmetry mates contributing to the binding site were correctly retained. Cryo-EM structures were handled as P1 crystal structures.

Systems containing unknown or metallo-organic residues were not retained, as these cannot be parameterized with AMBER ff19SB [15] or OpenFF 2.2.0 [16]. These force fields are required for the constrained energy-minimization step described below, which relaxes rebuilt missing loops and residues into a physically reasonable geometry and resolves minor inconsistencies in the binding sites; a system that cannot be parameterized cannot undergo this step. Three metallocofactors (heme (HEM), molybdopterin guanine dinucleotide (MGD), and the [4Fe-4S] cluster (SF4)) were exempted from this filter, as validated AMBER parameters are available for these specific residues [17, 18]. Other ligands were required to consist only of C, N, O, S, P, F, Cl, Br, and I, the only elements parameterized in OpenFF 2.2.0.

### Step 3: Structure repair

Non-standard amino acids and nucleotides, covalent modifications, and missing atoms were addressed using PDBFixer [19]. Systems containing any of these “defects” within the binding pocket (defined here as the 6 Å shell surrounding the ligand chain) were rejected to preserve the experimental integrity of the binding site. Outside the pocket, non-standard amino acids were substituted with their closest standard equivalents (e.g., hydroxyproline to proline, phosphoserine to serine). A custom rule was added to convert selenocysteine to cysteine, a substitution not handled by PDBFixer by default; residues without a predefined mapping were mutated to alanine. Non-standard nucleotides were mutated to adenosine for RNA chains and deoxy-adenosine for DNA chains. These substitutions were made, first of all because non-standard residues lack parameters in the AMBER force fields used later for energy minimization, but also because many downstream model architectures are not designed to handle such modifications.

Missing atoms and residues outside the pocket were rebuilt using PDBFixer’s default modeling routines, while covalent modifications outside the pocket were removed. Missing residues at chain termini were rebuilt up to a maximum of three residues. Water residues within the pocket were retained, as they may contribute to the interaction; outside the pocket, they were removed.

### Step 4: Protein-Ligand interaction quality control

PLI systems with at least one protein chain carrying a UniProtKB/Swiss-Prot annotation [20] were retained for further quality control.

Agreement between the atomic model and the underlying experimental data was quantified for each retained system but not used to exclude any complex. For structures determined by X-ray crystallography, we computed two complementary density-fit metrics [21, 22]: the real-space R-value (RSR), which quantifies the per-residue residual between observed and model-derived electron density, and the real-space correlation coefficient (RSCC), which measures the Pearson correlation between observed and calculated density maps over the residue volume. For structures determined by cryo-EM, where the experimental map is a reconstructed Coulomb potential map rather than a crystallographic electron density map, we instead computed the Q-score [23], which quantifies per-atom resolvability by comparing the map values in the neighborhood of each atom to a Gaussian reference. All three metrics were computed and reported separately for ligand and pocket residues.

The quality of the interaction itself was then evaluated using two geometric criteria. At least 10 interatomic close contacts between protein and ligand (atom pairs within 4 Å) were required, and the fraction of buried surface area of the ligand was required to exceed 40%. Solvent-accessible surface area was calculated using FreeSASA [24]. Jointly, these thresholds ensure that the ligand is engaged in a well-defined binding pose rather than merely adjacent to the protein surface.

### Step 5: Constrained energy minimization

To prepare the retained systems for minimization, all organic chains with fewer than 100 heavy atoms were first extracted as RDKit molecules [25], with bonds and bond orders assigned from the Chemical Component Dictionary (CCD) residue definitions [14]; inter-residue bonds were set to single bonds. The resulting molecules were protonated using Dimorphite-DL [26] with the improved rule set introduced in HiQBind [4], which incorporates chemically informed constraints such as suppressing protonation of amines directly bonded to non-hydrogen, non-carbon atoms (e.g., in sulfonamides). Protein chains were protonated with PDBFixer using standard pK_*a*_-based rules. All protonation states were assigned at physiological pH (7.4).

The prepared components were then reassembled into full systems in OpenMM 8.4.0 [19]. Force field parameters were assigned as follows: ff19SB for protein residues [15], OL21 and OL3 for nucleic acids [27], OPC3 for water and ions [28], and OpenFF 2.2.0 [16] with Gasteiger partial charges [29] for ligands and non-metal-coordinating cofactors. For the three supported metallocofactors (HEM, MGD, and SF4), AMBER parameters were converted to OpenMM format, as OpenFF 2.2.0 does not natively support metal-coordinating bonds.

Energy minimization adjusts atomic coordinates to lower the system’s force-field potential energy, following the energy gradient downhill toward the nearest local minimum to relieve steric clashes while leaving the overall fold intact. Minimization was performed using OpenMM’s L-BFGS implementation for up to 5,000 iterations [19], using a non-periodic topology and a 10 Å cutoff for non-bonded interactions. A restraint protocol was designed to relieve steric strain in the experimental structures while preserving close agreement with the original crystallographic geometry. Three different protocols were used for missing atoms (hydrogens and rebuilt atoms), pocket heavy atoms, and heavy atoms outside the pocket. Hydrogens and atoms rebuilt with PDBFixer were given no positional restraints, allowing free optimization. Other heavy atoms outside the pocket were subjected to a stiff harmonic positional restraint with energy *E*_*pos*_ = *kr*^2^, with *k* = 10^6^ kJ · mol^−1^ · nm^−2^, effectively freezing the protein scaffold. Heavy atoms within the pocket were instead restrained using a softer, flat-bottomed tethering potential (Eq. 1, Figure S1) [8, 30]:

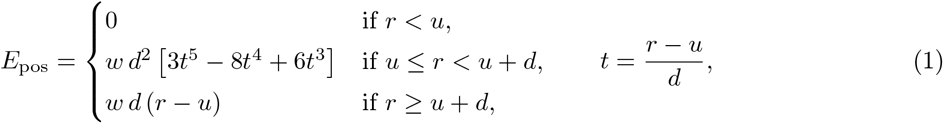

where *r* is the displacement from the crystallographic position, *u* = 0.25 Å is the flat-bottom width, *d* = 1.0 Å is the width of the smoothing region, and *w* = 10 kcal · mol^−1^ · Å^−2^. The flat-bottomed region permits atoms to move freely within the inherent coordinate uncertainty of typical crystallographic resolutions without energetic penalty. Beyond this tolerance, a quintic polynomial smoothly ramps up the restraint, which eventually becomes linear to avoid excessive penalties on displaced atoms. The transition polynomial in Eq. 1 is a *C*^2^-continuous quintic Hermite polynomial [31] satisfying Equation 2

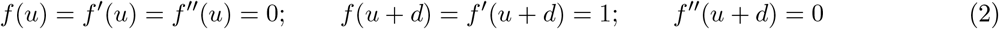

This construction guarantees continuity of the potential energy, force, and force gradient across all three regimes, a prerequisite for numerical stability and convergence of the minimizer.

### Hierarchical clustering

To support leakage-controlled train-test splitting for machine learning applications we provide cluster labels based on protein sequence, pocket sequence, ligand and interaction similarity as part of CROWN’s metadata annotation.

#### Protein sequence similarity

Amino acid sequences for all protein chains in CROWN are extracted from the processed structures and compared in an all-versus-all search with MMseqs2 [32]. From the resulting pairwise similarities, we construct a greedy chain mapping for every pair of CROWN complexes.

Pairwise sequence identities between all protein chains were computed with MMseqs2. Sequences were collected into a single database and searched all-versus-all with the database as both query and target, using an E-value cutoff of 0.01, a minimum sequence identity of 0.2, and a maximum of 5000 results per query; remaining parameters were left at their defaults. Fractional sequence identities (fident) were exported with convertalis and define *S*_*i*,*j*_in Eq. (1).

For a complex pair (*a, b*), let *a* denote the complex with the fewer chains and let *M*_*a*,*b*_ be the set of mapped chain pairs (*i, j*) with *i* ∈ *a* and *j* ∈ *b*. Global similarity is the length-weighted mean of the MMseqs2 chain similarities *S*_*i*,*j*_ (Eq. 3),

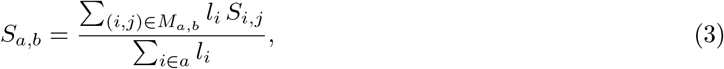

where *l*_*i*_ is the length of query chain *i*. Normalizing by the total length of all chains in *a*, rather than only the mapped ones, assigns a similarity of zero to any unmapped chain.

#### Pocket sequence similarity

The pocket (binding-site) residues of a CROWN entry is defined as all residues with at least one heavy-atom contact (*<* 4 Å) to the system ligand; we write *B*_*a*_ for the pocket residue set of entry *a*.

Pocket-level similarities are computed from a Smith–Waterman alignment [33] of the mapped chains, performed with Biopython [34]. For a residue *r* ∈ *B*_*a*_, let *π*_*a*→*b*_(*r*) denote the residue of *b* aligned to *r* under this alignment (undefined where *r* has no aligned partner). The fraction of shared pocket residues is the fraction of *B*_*a*_ whose aligned partner lies in *B*_*b*_ (Eq. 4),

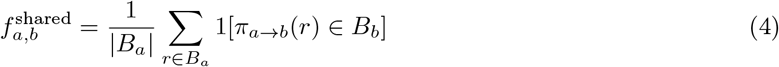

#### Ligand similarity

Pairwise ligand similarity was calculated as the Tanimoto similarity of ECFP4 fingerprints using RDKit [25, 35].

#### Interaction similarity

Protein–ligand interaction similarity was calculated using 32,768-bit PLEC fingerprints, which allow alignment-free comparison of complexes [36].

From these different similarity metrics, minimum spanning trees were created which can be used for single linkage clustering at any desired similarity cutoff.

### CASF-2016 benchmark

To assess the value of CROWN as a training resource, we benchmarked models trained on it against the CASF-2016 test set. CASF-2016 comprises 285 protein–ligand complexes grouped into 57 protein clusters and evaluates four complementary aspects of a scoring function: scoring, ranking, docking, and screening power [37]. The benchmark provides pre-generated docking poses, ensuring reproducibility.

We trained two knowledge-based scoring functions, DESPOT and DESPOT-screen (described in our companion manuscript [38]), on each of three training sets: raw CROWN structures (CROWN-xtal), energy-minimized CROWN structures (CROWN), and HiQBind [4]. To prevent train–test leakage, we excluded any training entry that both shared *>*70% sequence identity with a CASF-2016 protein chain and had *>*70% ECFP4 Tanimoto similarity to the associated CASF-2016 ligand.

Confidence intervals for all performance metrics were obtained by bootstrapping the set of test complexes over 10,000 resampling iterations, recomputing each metric on every replica and applying the bias-corrected and accelerated (BCa) interval estimator [**efron_better_1987**, 37]. To compare scoring functions against one another, we applied the non-parametric Friedman–Nemenyi test and treated two functions as statistically indistinguishable when *p >* 0.10. Both procedures follow the protocol of Su et al. [37], to which we refer for a full description of the performance tests.

Full details of the DESPOT scoring functions and their CASF-2016 evaluation are provided in the companion paper [38].

### Computational Details

All computations were performed on a Linux workstation equipped with an AMD Ryzen Threadripper PRO 5995WX processor (64 cores, 128 threads) and 252 GB of RAM. Generating the full CROWN dataset required approximately one week of wall-clock time.

## Results and Discussion

### Dataset Scope and Diversity

A direct comparison with established protein–ligand databases places CROWN in a distinctive position (Table 1). With 178,263 entries spanning 76,446 unique PDB identifiers and 18,124 unique UniProt accessions, CROWN represents an approximate four-fold increase in protein diversity relative to PDBBind [9] (3,627 UniProt IDs) and HiQBind [4] (2,705 UniProt IDs). Its ligand diversity is comparably broad: 31,005 unique CCD identifiers and 18,317 unique Murcko scaffolds [39], roughly doubling the chemical coverage of PDBBind and HiQBind.

**Table 1:** Comparison of dataset scope, ligand diversity, and ligand quality filtering across five structural protein–ligand interaction datasets. Values for CROWN are reported for the corrected structures.

| Property | PDBBind | HiQBind | BioLiP2 | PLINDER | CROWN |
| --- | --- | --- | --- | --- | --- |
| ***Dataset scope*** |  |  |  |  |  |
| Total entries | 19 449 | 31 573 | 86 458 | 649 915 | 178 263 |
| Unique PDB-CCD pairs | 17 758 | 17 247 | 22 720 | 201 836 | 93 347 |
| Unique PDB IDs | 17 758 | 17 088 | 17 701 | 111 867 | 76 446 |
| Unique UniProt IDs | 3 627 | 2 705 | 15 592 | 24 070 | 18 124 |
| Unique CATH IDs | 978 | 800 | 2 151 | 2 858 | 2 206 |
| Unique species | 488 | 395 | 1 438 | 1 991 | 1 699 |
| Affinity annotations | ✓ | ✓ | ✗ | ✗ | ✗ |
| ***Ligand diversity*** |  |  |  |  |  |
| Unique CCD IDs | 13 956 | 12 428 | 6 519 | 47 300 | 34 912 |
| Unique Murcko scaffolds | 8 251 | 7 660 | 3 213 | 22 743 | 18 317 |
| Ion ligands | 0 | 0 | 1 805 | 22 728 | 0 |
| Covalent ligands | 870 | 24 | 1 708 | 32 276 | 0 |
| Artifact ligands | 15 | 34 | 523 | 18 626 | 0 |
| ***Structure issues*** |  |  |  |  |  |
| Missing bonds | 990 | 114 | 2 104 | 37 422 | 0 |
| Steric overlaps | 323 | 313 | 656 | 5 670 | 0 |
| Unresolved ligand atoms | 463 | 1 | 1 564 | 18 815 | 0 |
| Unresolved pocket atoms | 1 102 | 955 | 1 579 | 12 457 | 0 |
| Non-standard pocket residues | 307 | 245 | 972 | 4 473 | 0 |
| ***Structure corrections*** |  |  |  |  |  |
| Protonation | ✓ | ✓ | ✗ | ✗ | ✓ |
| Energy minimization | ✗ | ✗ | ✗ | ✗ | ✓ |

The comparison with BioLiP2 [40] is instructive. Although BioLiP2 catalogs 86,458 entries and achieves high species diversity, its ligand diversity is markedly lower than that of CROWN: only 6,519 unique CCD identifiers and 3,213 Murcko scaffolds. This discrepancy likely reflects BioLiP2’s inclusion of ions, artifacts, and other non-drug-like entities that inflate entry counts without expanding the chemical space relevant to drug discovery and molecular recognition. By explicitly removing these categories, CROWN achieves a cleaner and more chemically focused dataset.

### Ligand Property Distributions

The distribution of molecular properties across CROWN’s ligand set differs meaningfully from that of existing databases (Figure S2). Compared with CROWN, PDBBind — for years the dominant training resource for scoring-function development — markedly under-represents bulkier ligands: very few entries exceed 50 heavy atoms or 12 hydrogen-bond acceptors. CROWN instead preserves the broader molecular-weight distribution characteristic of PLINDER, including the larger, more complex ligands increasingly relevant to contemporary drug discovery, such as PROTACs [41], macrocycles, and molecular glues, while removing the ions, solvents, and small fragments that skew PLINDER’s lower end. The resulting heavy-atom-count distribution is therefore both broader and cleaner, better representing the diversity of therapeutically relevant small molecules. Similar trends hold for rotatable-bond counts and for hydrogen-bond acceptor and donor counts, where CROWN provides broader coverage than either PDBBind or HiQBind. This extended coverage matters for training models expected to generalize across ligand sizes and polarities, since models trained on narrow property distributions tend to perform poorly on out-of-distribution compounds. The distribution of quantitative estimate of drug-likeness (QED) [42] values in CROWN closely mirrors those of BioLiP2 and PLINDER, spanning a wide range of drug-likeness.

### Structure Quality and Annotation Completeness

This completeness extends to atomic-level and geometric quality. CROWN contains zero entries with unresolved ligand atoms, missing bonds, steric clashes, or non-standard residues in the binding pocket (Table 1). By contrast, PDBBind retains 463 entries with unresolved ligand atoms, 990 with missing bonds, and 323 with steric overlaps, while PLINDER contains more than 18,000 entries with unresolved ligand atoms, over 37,000 with missing bonds, and over 5,600 with steric clashes. Individually minor, such inconsistencies collectively distort learned representations of interaction patterns.

### Representative Structural Corrections

Figure 2 illustrates several representative issues automatically detected and corrected by the pipeline. Figure 2a (PDB ID: 3FIV) shows how alternate conformer occupancies create an ambiguous atomic representation, which the CROWN pipeline resolves by selecting the highest-occupancy conformer. In PDB entry 8GRT (Figure 2b), missing inter-residue covalent bonds produce a fragmented chain representation; the pipeline detects these gaps and repairs the connectivity into a single continuous chain. In Figure 2c (PDB ID 5CHR), an oxygen cofactor chain exhibits severe steric clashes with the system ligand, and the pipeline removes the offending chain to preserve a physically reasonable binding site. Finally, the PDB entry for 6ARO (Figure 2d) contains only a single protein chain in the ligand-binding site, whereas symmetry-mate expansion reveals a second chain in close contact on the opposite side of the ligand. Though individually straightforward, these corrections (absent from popular databases such as PLINDER) are pervasive across the PDB and would be prohibitively labor-intensive to address manually at CROWN’s scale.

**Figure 2:**
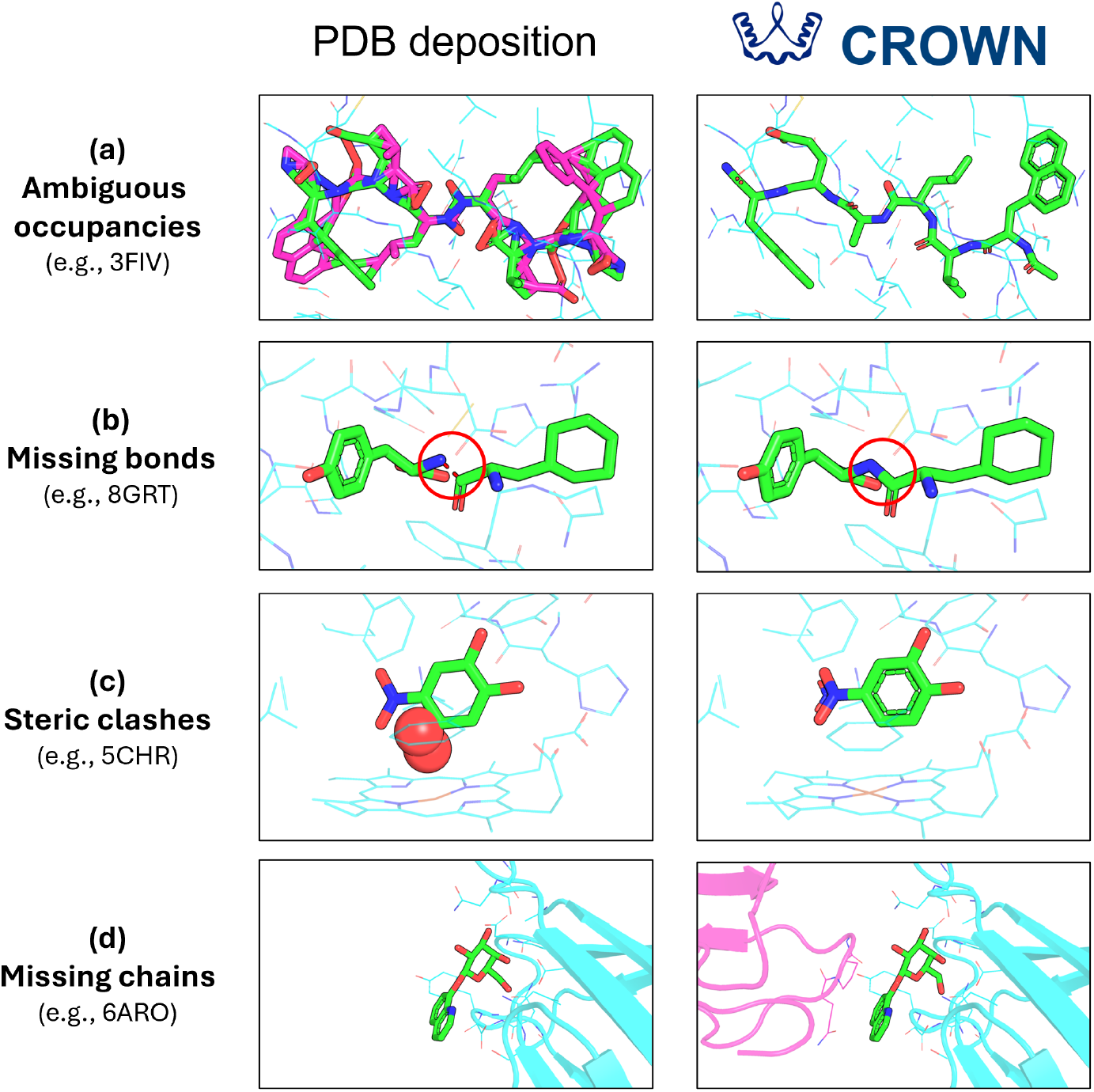
Representative structural corrections applied by the CROWN pipeline, shown before (left) and after (right) processing. (a) Ambiguous alternate-conformer occupancies in PDB 3FIV are resolved by retaining the highest-occupancy conformer. (b) Missing inter-residue covalent bonds in PDB 8GRT are detected and repaired, merging disconnected residues into a single continuous chain. (c) A cofactor chain clashing severely with the system ligand in PDB 5CHR is identified and removed. (d) Symmetry-mate expansion recovers a second binding-site chain absent from the deposited PDB entry for 6ARO.

### Constrained energy minimization

Experimental structures are not free of geometric artifacts: steric clashes and strained bond geometries are common, and refinement practices vary widely across PDB depositions. Moreover, the rebuilt missing atoms and residues introduced in the structure repair step of the CROWN pipeline also require energy minimization for correct placement. Left uncorrected, such inconsistencies propagate directly into any model trained on the coordinates, and a network has no way to distinguish a genuine interaction geometry from an artifact. By relaxing each structure toward a physically reasonable local energy minimum while restraining it close to the deposited coordinates, minimization removes such artifacts without erasing the experimental signal. For structure-based machine learning this yields a more uniform training target: clashes, distorted geometries, and depositor-to-depositor variation are reduced, so that models learn the physics of molecular recognition rather than artifacts specific to individual source structures.

Figure 3a confirms that the tiered restraint strategy performs as intended across the structural components of each complex. The protein scaffold, under stiff harmonic restraints, shows negligible displacement, while pocket and ligand heavy atoms, governed by the flat-bottomed potential with its 0.25 Å tolerance, exhibit median RMSD values of approximately 0.22 and 0.27 Å, respectively. These displacements fall pre-dominantly within or just above the flat-bottom width, indicating that most atomic adjustments are comparable in magnitude to the coordinate uncertainty of typical medium-resolution crystal structures [7], with only modest additional movement where strain relief is required. The clear separation between the RMSD distributions demonstrates that the restraint hierarchy localizes structural adjustments to the binding site while preserving the overall crystallographic geometry.

**Figure 3:**
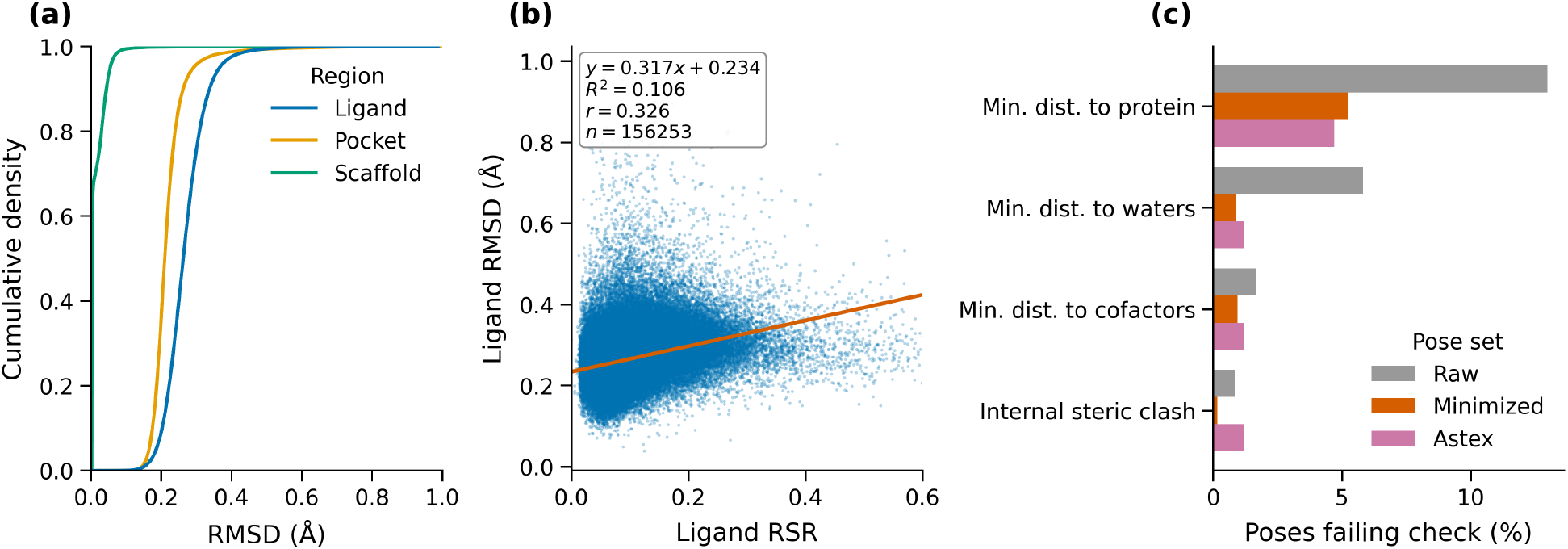
Effects of constrained energy minimization. **a)** Empirical cumulative distribution functions of the RMSD between crystallographic and energy-minimized coordinates, separated by ligand, pocket and scaffold regions. **b)** Scatterplot indicating the correlation between ligand real-space R-value (RSR) and the RMSD between crystallographic and energy-minimized coordinates. **c)** Failure rates of common Posebusters checks, compared between the raw CROWN structures, energy-minimized CROWN structures and the Astex Diverse Set.

Figure 3b shows a positive correlation between ligand RSR and the RMSD between crystallographic and minimized coordinates (for the subset of CROWN where RSR values are available). Although the relationship is too weak to be predictive at the level of individual ligands, it confirms the expected trend: more correction is required for poorly resolved ligands.

Finally, figure 3c reports failure rates for four common PoseBusters checks [43], comparing crystallographic and energy-minimized poses. Constrained energy minimization substantially reduces these rates: for example, clashes with protein chains fall from 12.97% to 5.23%, clashes with surrounding waters from 5.83% to 0.89%, and internal steric clashes from 0.85% to 0.17%. When compared to the PoseBusters statistics of the highly curated Astex Diverse Set [3], we see that the receptor-ligand steric clash rates of the energy-minimized CROWN entries are very similar to those of the Astex set.

Together, the above results confirm that CROWN’s restrained energy minimization significantly reduces structural artifacts while maintaining the overall geometry.

### Multi-axis clustering for leakage-controlled data splits

A recurring problem in structure-based machine learning is that protein–ligand databases are highly redundant, with near-identical complexes represented across many PDB entries. Under a random train–test split such redundancy leaks across the partition, so a model is effectively evaluated on variants of structures it has already seen, and reported performance overstates true generalization. Grouping related entries and splitting along cluster boundaries removes this leakage and yields more honest estimates of out-of-distribution performance.

We therefore provide cluster labels for every CROWN entry along three axes — protein sequence, ligand chemistry, and interaction fingerprint (PLEC) — because redundancy can enter through each independently: two complexes may share a protein, a common ligand, or a similar interaction pattern without sharing the other two. Offering all three lets users control the generalization axis a given task requires — novel proteins, novel ligands, or novel interaction modes — and combine them for stricter splits.

Figure S3a demonstrates how the cluster-size distributions in CROWN are strongly skewed across all three schemes, comprising many small clusters and a few large ones. For protein clusters, 80% of the data lies in 22.9% of the clusters; for ligand clusters, 80% lies in 13.3% of the clusters and 50% within just 0.2%; interaction clustering is the least skewed, with 80% of the data in 49.7% of the clusters. This imbalance bears directly on train-test splitting strategies: because a few dominant clusters carry a large share of the data, split assignments must account for cluster size to avoid placing a disproportionate fraction of the dataset on one side.

We next asked how much information the three schemes share (Figure S3b). Agreement with the Adjusted Mutual Information (AMI) was quantified — the appropriate chance-corrected measure when clusterings are unbalanced and contain many small clusters [44], precisely the regime established in Figure **??**a. AMI measures how much knowing an entry’s label under one scheme reduces uncertainty about its label under another, on a scale where independent partitions score 0 and identical partitions score 1. All three schemes share appreciable information, confirming that they are not independent. Protein sequence and PLEC clusters are the most strongly related, with an AMI of 56.9%.

Together these results show that the three clusterings are related but not redundant: each captures distinct structure in the data, so combining them supports more rigorous train–test splits than any single axis alone.

### Web Interface

The complete CROWN dataset is freely accessible through a dedicated web interface at https://crown.lbmd.be (Figure 4), which offers two primary modes of interaction: search and browse. The search module supports advanced multi-field filtering and sorting, allowing users to query entries by structural quality metric (e.g., X-ray resolution, RSR, RSCC), ligand property (e.g., CCD code, molecular weight), or biological annotation (e.g., UniProt ID, NCBI Taxon ID, CATH classification). The browse view presents a detailed summary for each entry, organizing protein and ligand metadata alongside an interactive 3D structure viewer. Users can inspect both the corrected crystal coordinates prior to energy minimization and the final energy-minimized structures, and can superimpose the two conformations to assess the extent of structural rearrangement introduced during refinement. Beyond the web interface, the full dataset is available for bulk download: receptors (including any cofactors) are provided as PDB files and system ligands as SDF files, with both crystal and energy-minimized coordinates included. A similarity-search feature built on the clustering schemes above enables retrieval of entries with related binding-site features.

**Figure 4:**
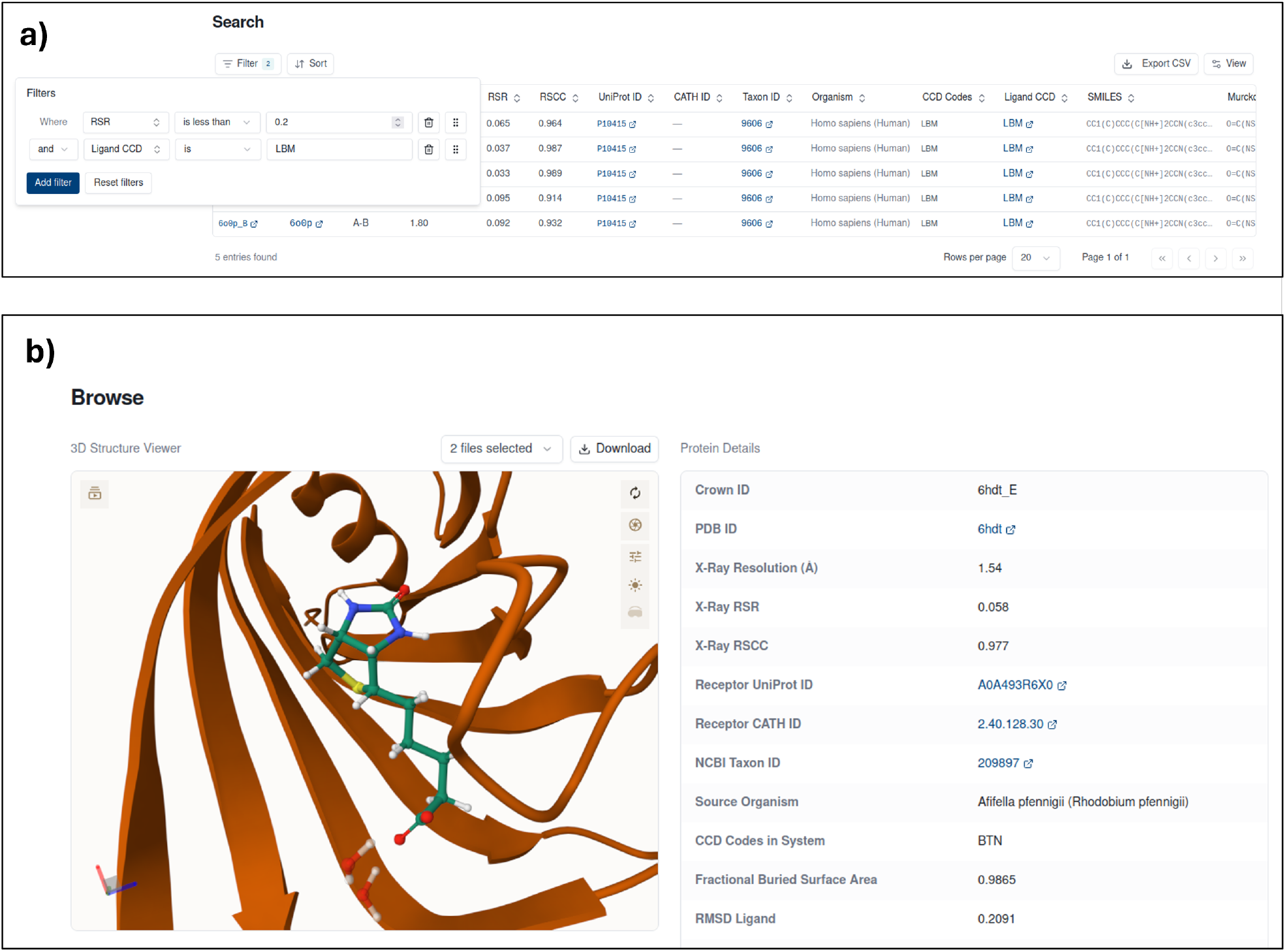
Overview of the CROWN web interface. **a)** The search view, supporting advanced filtering and sorting across multiple structural, chemical, and biological annotation fields. Here, an example query is shown filtering for entries with an RSR below 0.2 and ligand CCD code LBM. **b)** The browse view for an individual entry (PDB: 6hdt), displaying an interactive 3D structure viewer alongside protein metadata (resolution, quality metrics, organism, receptor classification) and ligand details (SMILES, Murcko scaffold, molecular weight, rotatable bonds). Both the crystal and energy-minimized coordinates can be visualized and compared within the viewer.

### CROWN as a training resource

Training DESPOT and DESPOT-screen on each of the three training sets yielded six models, which we evaluated on the full CASF-2016 benchmark (Figure 5).

**Figure 5:**
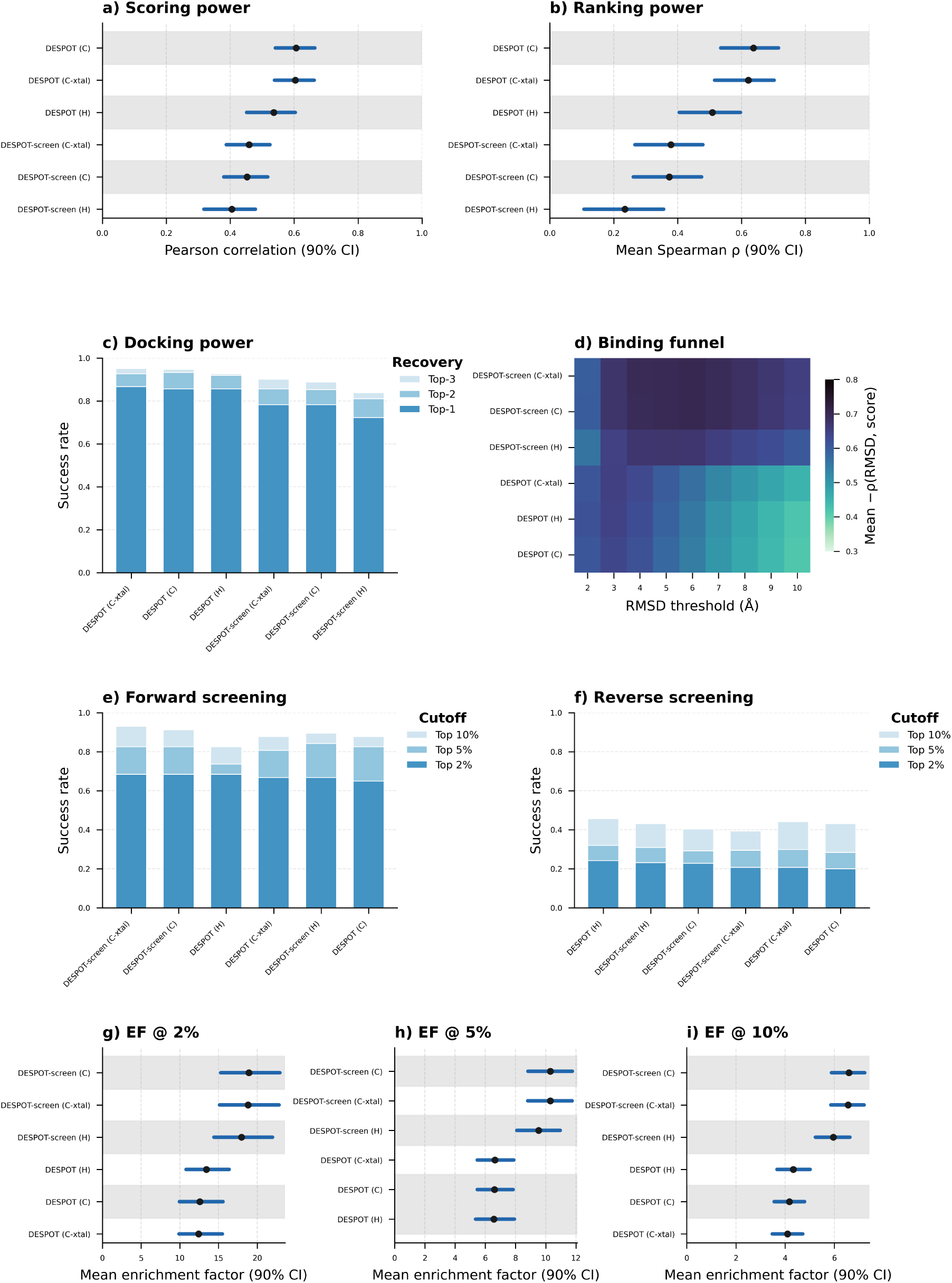
CASF-2016 benchmark performance compared between CROWN and HiQBind training sets. **a)** Scoring power: Pearson correlation between predicted scores and experimental binding affinities. **b)** Ranking power: mean Spearman correlation for ranking ligands within protein targets. Black dots indicate point estimates; blue bars indicate 90% bootstrap confidence intervals. Scoring functions on the same white/gray band are statistically indistinguishable (Friedman-Nemenyi test, p *>* 0.10). **c)** Docking power: success rate for identifying near-native poses (RMSD *<* 2.0 Å) at top-1, top-2, and top-3 rankings. **d)** Binding funnel analysis: Spearman correlation between score and RMSD as a function of RMSD threshold. **(e-f)** Screening power: success rate for forward screening (identifying correct ligands for targets) and reverse screening (identifying correct targets for ligands) at various ranking thresholds. **(g-i)** Enrichment factors at 21%, 52%, and 105% thresholds. The training set choice is indicated in parentheses as (C) for CROWN, (C-xtal) for the raw CROWN structures or (H) for HiQBIND.

Scoring and ranking power both improved when models were trained on CROWN rather than HiQBind (Figure 5a–b). The effect on ranking power was most pronounced: DESPOT’s mean Spearman correlation between score and affinity rose from 0.509 (HiQBind) to 0.637 (CROWN), and DESPOT-screen’s from 0.235 to 0.374. The Pearson correlation between score and affinity rose from 0.536 to 0.606 for DESPOT and from 0.404 to 0.452 for DESPOT-screen.

The docking-power results were driven mainly by DESPOT-screen. For the HiQBind-trained model, the Spearman correlation between score and RMSD to the crystal pose dropped markedly across the binding funnel (Figure 5d), which likely contributed to its lower pose recovery: top-1 recovery fell from 0.789 (CROWN) to 0.719 (HiQBind) (Figure 5c).

Screening power was least sensitive to the choice of training set, although DESPOT-screen’s increase in enrichment factors for the CROWN-trained models was still statistically significant (Figure 5g–i).

Together, these results show that training on CROWN significantly improves model performance, most clearly in scoring, ranking, and docking power. The gap between the two CROWN variants was much smaller: differences between energy-minimized CROWN and raw CROWN (CROWN-xtal), while statistically significant, were minor. For this benchmark—where models see only the binding site rather than the full protein—the two sets are therefore largely interchangeable. This does not extend to every application: in tasks such as ligand-conditioned protein design [10], newly built loops and residues distant from the binding site must be properly minimized. We therefore release both variants but recommend the energy-minimized structures for the majority of use cases.

### Pipeline Development: From Manual Curation to Automation

Starting from 201,124 experimental structures, the pipeline yields 178,263 protein–ligand complexes. Of the 258,241 candidate complexes initially defined, 22.48% fail during structure repair owing to gaps, missing atoms, or non-standard residues in the binding site; of the remainder, 89.51% pass all quality-control filters, and 99.48% of those undergo successful constrained energy minimization. The high pass rate of the final stage reflects not the simplicity of the problem but the cumulative effort invested in resolving the long tail of structural edge cases that characterize real-world PDB data (see Methods and code documentation). Because the finished pipeline is fully automated, it can be rerun end-to-end as new structures are deposited; we will use this to release annual updates of CROWN that track the continued growth of the PDB.

The effort behind CROWN’s high pass rates points to a distinction worth making explicit: between the automation of the finished pipeline and the manual, iterative process required to design it. The structural heterogeneity of the PDB is immense, deposited structures vary widely in their conventions for alternate conformers, chain annotations, bond connectivity, residue naming and crystallization artifacts, and no single set of rules can be written *a priori* to handle it. The automated pipeline instead emerged from repeated rounds of testing against large batches of structures, manual inspection of failures, and incremental rule refinement until pass rates stabilized. We stress this because it bears directly on reproducibility and on future work: the apparent simplicity of an automated pipeline can obscure the expert curation beneath it, and extending CROWN to new structure types (e.g., NMR) or chemical categories (e.g., covalent inhibitors, metallodrugs) will likely require a comparably intensive effort.

### Current Limitations

Despite its strengths, the current CROWN pipeline still has several limitations. The most significant arises from force-field coverage. OpenFF 2.2.0 [16], used to parametrize ligands and non-metal-coordinating cofactors, does not support metal-coordination bonds or ligands containing rare elements such as boron, selenium, and silicon. Although we developed an AMBER-derived workaround for three common metal-coordinating cofactors (HEM, MGD, and SF4), this solution is inherently limited and cannot be generalized to the full diversity of metalloprotein–ligand interactions. A small number of PLI systems also involve bonding patterns that fall outside the current coverage of OpenFF 2.2.0 and were not retained.

A related limitation concerns non-standard amino acids. AMBER ff19SB [15] provides parameters for the 20 canonical residues but not for post-translationally modified or otherwise non-standard amino acids. Outside the binding pocket, these residues are substituted with their closest standard equivalents, a reasonable approximation given their distance from the interaction of interest. Within the pocket, however, such substitutions could compromise the fidelity of the local interaction environment, so these entries were removed.

### Future Directions

The choice of structural dataset should be guided by the intended machine learning task, as no single resource is optimal for all applications. Models that rely on experimentally measured binding affinities are best trained and evaluated using datasets such as PDBBind [9] and HiQBind [4], while applications centered on for example metal coordination are better served by broader databases such as PLINDER [5]. For most other structure-based learning tasks involving drug-like protein–ligand complexes, we propose CROWN as a large, uniformly processed dataset that can serve as a general-purpose foundation for model training, benchmarking, and method development.

In future, we hope to further extend CROWN’s coverage, for example by including chemistries currently beyond the reach of available open-source force fields. Because CROWN is generated directly from the PDB, such extensions would integrate naturally into a framework that supports automatic, reproducible dataset updates as the PDB grows.

## Conclusion

We have presented CROWN, a dataset of 178,263 curated protein–ligand complexes produced by a fully automated preprocessing pipeline. By combining stringent quality filtering with structural correction, protonation under physiological pH, and constrained energy minimization, CROWN offers roughly four-fold greater protein diversity than existing curated resources, broader ligand chemical-space coverage, and complete electron-density quality annotations. Its geometry-centric design encompasses the full breadth of well-resolved PDB structures regardless of whether affinity measurements have been reported. Thus, CROWN can serve as a training and benchmarking resource for structure-based machine learning — including generative pose models, structure prediction, ligand-conditioned sequence design, and scoring-function development — without depending on scarce affinity labels.

We validated this premise on the CASF-2016 benchmark, where knowledge-based scoring functions trained on CROWN outperformed the same models trained on HiQBind across scoring, ranking, and docking power, while the energy-minimized and raw CROWN variants proved largely interchangeable for this binding-site–only task. This confirms that CROWN’s scale and diversity translate into concrete downstream gains rather than merely broader coverage.

Central to the pipeline is the constrained energy-minimization step (absent from all widely used protein–ligand datasets), which reconciles the heterogeneous refinement practices across PDB depositions into a structurally uniform collection. Because the entire pipeline is fully automated, it can be rerun as the PDB grows; we will release annual updates to keep the dataset current with newly deposited structures. We expect that CROWN and its preprocessing framework will provide a durable foundation for next-generation structure-based models of molecular recognition.

## Abbreviations

PLI: protein-ligand interaction
PDB: Protein Data Bank
mmCIF: macromolecular Crystallographic Information File
CCD: Chemical Component Dictionary
RSR: real-space R-value
RSCC: real-space correlation coefficient
RMSD: root-mean-square deviation
QED: quantitative estimate of drug-likeness
ECFP: extended-connectivity fingerprint
PLEC: protein-ligand extended connectivity
AMI: adjusted mutual information
SASA: solvent-accessible surface area
PTM: post-translational modification
PROTAC: proteolysis-targeting chimera
L-BFGS: limited-memory Broyden–Fletcher–Goldfarb–Shanno

## Data and Software Availability

The complete CROWN dataset is freely available at https://crown.lbmd.be under a CC-BY 4.0 license. For each of the 178,263 protein–ligand complexes, protonated receptor structures are provided as PDB files and ligands together with any associated cofactors as SDF files. Both the corrected crystal coordinates prior to energy minimization and the final energy-minimized coordinates are included. Each entry is accompanied by metadata covering crystallographic quality metrics (resolution, RSR, RSCC), biological annotations (UniProt ID, CATH classification, source organism), and ligand descriptors (SMILES, Murcko scaffold, molecular weight).

The full dataset can be downloaded as a bulk archive at https://zenodo.org/records/20825315. The full preprocessing pipeline for CROWN is available as open-source software at https://github.com/KUL-LBMD/CROWN under the MIT license. The repository includes all pipeline stages described in this work — quality filtering, structural correction, protonation, force field assignment, constrained energy minimization, and post-minimization validation — along with documentation, environment specifications, and example usage. Dependencies include BioPython, RDKit, Dimorphite-DL and OpenMM 8.4.0; full version information is provided in the repository.

DESPOT is available as open-source software at https://github.com/KUL-LBMD/DESPOT under the MIT license. The repository includes scripts for training and inference with all DESPOT variants, along with documentation, environment specifications, and example usage. All pre-trained DESPOT models are available at https://zenodo.org/records/20829559

## Author Contributions

R.P.: Conceptualization, Methodology, Data Curation, Investigation, Formal Analysis, Software, Writing – Original Draft. W.V.E. & B.B. (Bruncsics, Balint): Software, Writing - review and editing. B.B. (Bruncsics, Bence) & A.A.: Methodology, Writing - review and editing. Y.M. & A.V.: Supervision, Conceptualization, Methodology, Writing - review and editing.

## Acknowledgments

Robin Poelmans acknowledges support from the Research Foundation - Flanders (FWO) through a PhD fellowship (Grant No. 1176225N). Arnout Voet and Bence Bruncsics acknowledge funding through the Citribel Research Chair in Protein Engineering for Circular Waste Processes. Adam Arany and Yves Moreau are affiliated with Leuven.AI and received funding from the Flemish Government (AI Research Program).

## Notes

### Competing Interest Statement

The authors have declared no competing interest.

### Summary of Updates

In this revised version, we have extended CROWN's scope to also include complexes without available RSR info, as well as complexes stemming from cryo-electron microscopy, thus extending CROWN from 141,261 to 178,263 curated structures. Furthermore, to evaluate CROWN as a training resource, a CASF-2016 benchmark evaluation is now included for the DESPOT and DESPOT-screen scoring functions trained on CROWN (Figure 5). The supplementary information is updated to give more information on the usage of CROWN and its associated metadata.

