## Supplementary Information for "CROWN: Curated Repository Of Well-resolved Non-covalent interactions"

### Supporting Information for: "CROWN: Curated Repository Of Well-resolved Non-covalent interactions"

#### Data Usage

The CROWN dataset can be found in the following Zenodo repository: `https://zenodo.org/records/20825315`, which contains the following files:

`crown.tar.gz` Protein–ligand complex structures in CROWN.

`CROWN_metadata.parquet` Metadata for the CROWN dataset (one row per complex).

`CROWN_combined_mst.parquet` Minimum spanning trees of the CROWN cluster metrics.

`README.md` Description of the structural dataset and the columns in the metadata file.

Unzipping the data tarball with `tar -xzvf crown.tar.gz` yields the directory `complexes/`, which contains one subdirectory per complex (named by PDB ID and binding chain label). For example, `complexes/3zwe_F/` contains:

`receptor.pdb` All protein and additive (water, ions, cofactors) chains within 4 Å of the ligand structure.

`receptor_minimized.pdb` The same chains after energy minimization.

`ligand.sdf` Protonated ligand structure.

`ligand_minimized.sdf` Protonated and energy-minimized ligand structure.

A description of all fields in the metadata file can be found in Table S1.

Table S1: Description of the fields in the CROWN test set. Each row is a protein–ligand complex; list-valued fields may contain multiple entries.

| Field | Type | Description | Example |
|---|---|---|---|
| basename | string | Unique entry identifier: PDB ID followed by the binding chain label(s). | 1w20_LA |
| MW | float | Ligand molecular weight (Da), computed from the SMILES. | 308.263 |
| HeavyAtoms | float | Number of non-hydrogen (heavy) atoms in the ligand. | 21.0 |
| N+O_Atoms | float | Combined count of nitrogen and oxygen atoms in the ligand. | 10.0 |
| HBD | float | Number of hydrogen-bond donors (Lipinski definition). | 6.0 |
| HBA | float | Number of hydrogen-bond acceptors (Lipinski definition). | 9.0 |
| RotatableBonds | float | Number of rotatable bonds in the ligand. | 5.0 |
| NumRings | float | Number of rings in the ligand. | 1.0 |
| TPSA | float | Topological polar surface area of the ligand ($Å^2$). | 179.61 |
| QED | float | Quantitative estimate of drug-likeness, in [0,1]. | 0.2891 |
| SMILES | string | Canonical SMILES string of the ligand. | CC(=O)N[C@H]1[C@H]([C@H](O)[C@H](O)CO)O[C@@](O)(C(=O)[O-])C[C@@H]1O |
| MurckoScaffold | string | Bemis–Murcko scaffold of the ligand, as SMILES. | C1CCOCC1 |
| pdb_id | string | Four-character PDB accession code of the source structure. | 1w20 |
| lig_name | string | Three-character PDB chemical-component (ligand) identifier. For oligomers, CCD codes are joined with '_' | SIA |
| sas_ratio | float | Fractional buried surface area of the ligand in the complex. | 0.990311 |
| ligand_rsr | float | Real-space R-factor (RSR) for the ligand; lower is better. | 0.102 |
| ligand_rscc | float | Real-space correlation coefficient (RSCC) for the ligand; higher is better. | 0.873 |
| pocket_rsr | float | Real-space R-factor for the binding-pocket residues. | 0.054292 |
| pocket_rscc | float | Real-space correlation coefficient for the binding-pocket residues. | 0.962542 |
| global_q | float | Q-score for the whole structure (Cryo-EM only). | 0.392 |
| ligand_q | float | Q-score for the ligand (Cryo-EM only). | 0.341 |
| pocket_q | float | Q-score for binding pocket residues (Cryo-EM only). | 0.420684 |
| chain_set | string | Hyphen-separated set of chain identifiers defining the assembly. | LA-D |
| resolution | float | Experimental resolution of the structure (Å). | 2.08 |
| r_free | float | Crystallographic $R_{free}$ of the deposited structure. | 0.195 |
| experimental_method | string | Structure-determination method (e.g. X-ray, Cryo-EM). | X-ray Crystallography |
| uniprot_id | list[string] | UniProt accession(s) of the target protein(s). | Q6XV27 |
| taxon_id | list[float] | NCBI taxonomy identifier(s) of the source organism(s). | 383550.0 |
| species_name | list[string] | Source organism name(s). | Influenza A virus (strain A/Duck/England/1/1956 H11N6) |

*continued on next page*

*Table S1 continued*

| Field | Type | Description | Example |
|---|---|---|---|
| `entry_name` | list[string] | UniProt entry name / mnemonic(s). | `NRAM_I56A2` |
| `GO_ids` | list[string] | Associated Gene Ontology term identifier(s). | `GO:0004308` |
| `protein_name` | list[string] | Descriptive protein name(s) from UniProt. | `Neuraminidase (EC 3.2.1.18)` |
| `EC_number` | list[string] | Enzyme Commission number(s), where applicable. | `3.2.1.18` |
| `cath_ids` | list[string] | CATH structural-classification identifier(s). | `2.120.10.10` |
| `Ligand_RMSD` | float | Ligand RMSD after energy minimization (Å).[†] | `0.3329` |
| `Pocket_RMSD` | float | RMSD of the binding-pocket atoms after energy minimization (Å). | `0.221` |
| `Scaffold_RMSD` | float | RMSD of protein scaffold (outside of pocket) after energy minimization (Å). | `0.0044` |
| `Rebuilt_RMSD` | float | RMSD of rebuilt residues and loops after energy minimization (Å). | `1.7366` |
| `H_RMSD` | float | RMSD of H atoms after energy minimization (Å). | `0.5419` |
| `0.5 ligsim cluster` | integer | Cluster ID from ligand-similarity clustering at a 0.5 threshold. | `5046` |
| `0.7 ligsim cluster` | integer | Cluster ID from ligand-similarity clustering at a 0.7 threshold. | `14707` |
| `0.9 ligsim cluster` | integer | Cluster ID from ligand-similarity clustering at a 0.9 threshold. | `21416` |
| `0.5 pli-sim cluster` | integer | Cluster ID from protein–ligand-interaction similarity at 0.5. | `2087` |
| `0.7 pli-sim cluster` | integer | Cluster ID from protein–ligand-interaction similarity at 0.7. | `2435` |
| `0.9 pli-sim cluster` | integer | Cluster ID from protein–ligand-interaction similarity at 0.9. | `2662` |
| `0.5 seq-sim cluster` | integer | Cluster ID from protein-sequence similarity at a 0.5 threshold. | `1` |
| `0.7 seq-sim cluster` | integer | Cluster ID from protein-sequence similarity at a 0.7 threshold. | `256` |
| `0.9 seq-sim cluster` | integer | Cluster ID from protein-sequence similarity at a 0.9 threshold. | `2088` |
| `0.5 pocketsim cluster` | integer | Cluster ID from binding-pocket similarity at a 0.5 threshold. | `128` |
| `0.7 pocketsim cluster` | integer | Cluster ID from binding-pocket similarity at a 0.7 threshold. | `166` |
| `0.9 pocketsim cluster` | integer | Cluster ID from binding-pocket similarity at a 0.9 threshold. | `255` |

For all four cluster metrics, single-linkage cluster labels are pre-computed in `CROWN_metadata.parquet` at cutoffs of 50%, 70%, and 90%. For custom thresholds, the minimum spanning trees (MSTs) in `CROWN_combined_mst.parquet` can be used instead. Each MST is stored as four columns:

`metric` Similarity metric: one of `seq-sim`, `pocket-sim`, `lig-sim`, or `pli-sim`.

`id1` CROWN ID of the first complex.

`id2` CROWN ID of the second complex.

`similarity` Pairwise similarity between `id1` and `id2` for the given metric.

A helper script for using the MSTs is provided in the CROWN GitHub repository (`https://github.com/KUL-LBMD/CROWN`). First ensure that `CROWN_metadata.parquet` and `CROWN_combined_mst.parquet` are present in `CROWN/data/metadata/`, then run, for example:

```
python scripts/make_clusters.py label --metric seq-sim --threshold 0.4
```

#### Supplementary Information

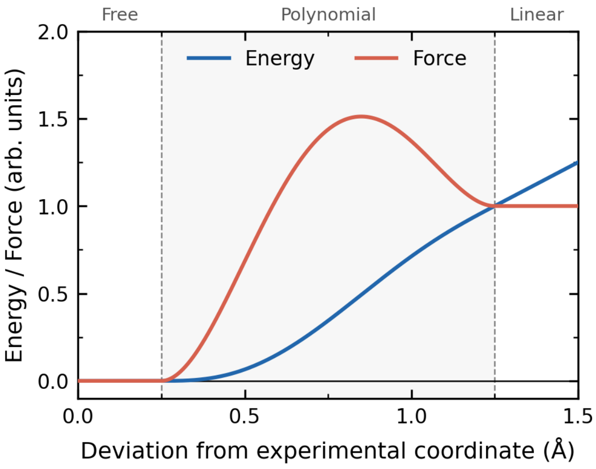


Figure S1: **Flat-bottomed tethering potential applied to heavy atoms within the pocket during constrained energy minimization.** The potential comprises three regimes: a force-free zone ($r < 0.25$ Å) accommodating coordinate uncertainty, a $C^2$-continuous quintic polynomial transition region that smoothly ramps up the restraint, and a linear regime preventing excessive penalties at larger displacements. Energy (blue) and force (red) profiles are shown as a function of deviation from the original position.

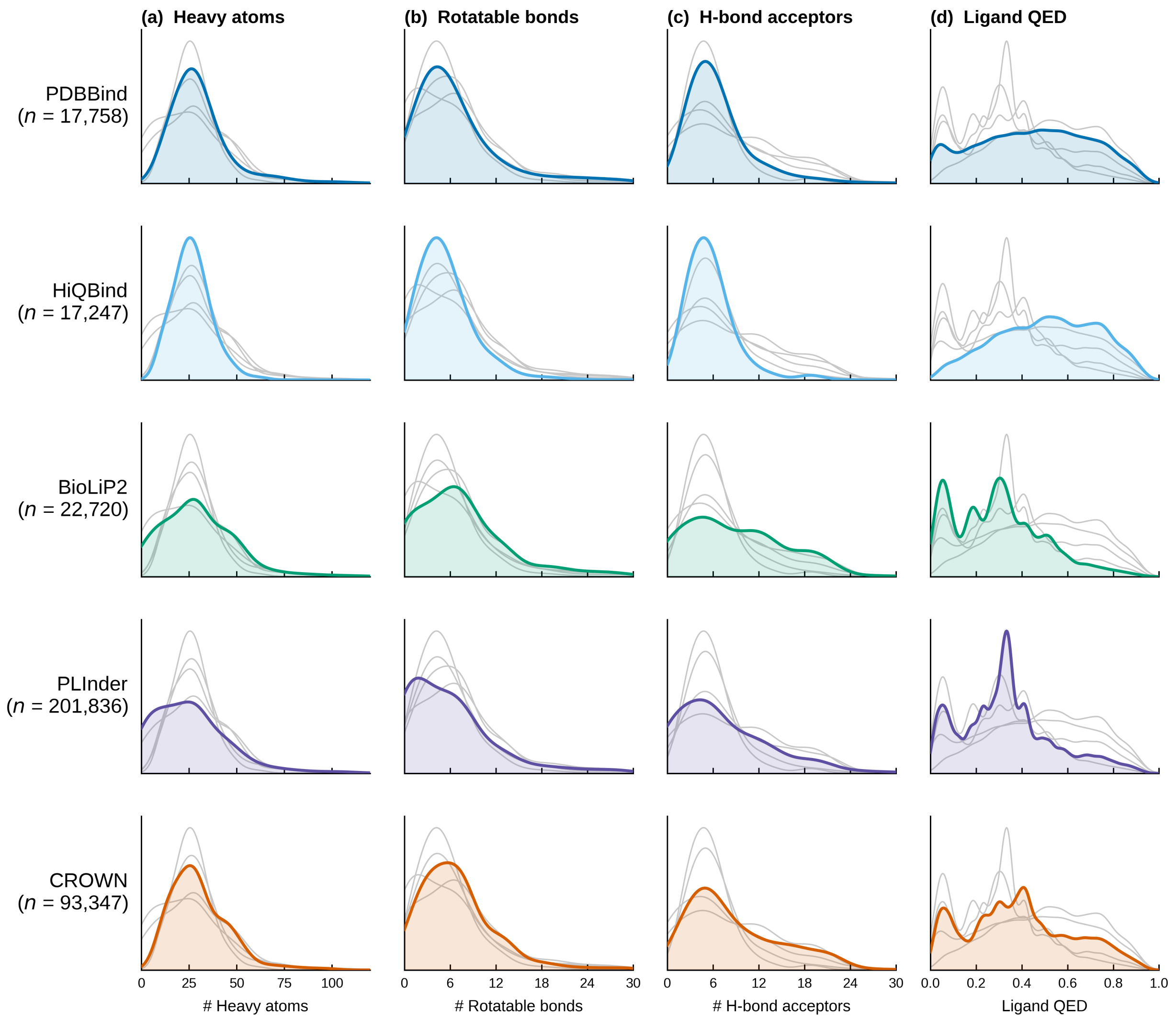


Figure S2: **Ligand property distributions across five protein–ligand interaction datasets** (PDB-Bind, HiQBind, BioLiP2, PLINDER, and CROWN). Kernel density estimates are shown for (a) heavy-atom count, (b) rotatable-bond count, (c) hydrogen-bond-acceptor count, and (d) ligand Quantitative Estimate of Drug-likeness (QED) score. All the density curves were computed over the unique PDB–CCD pairs in each dataset.

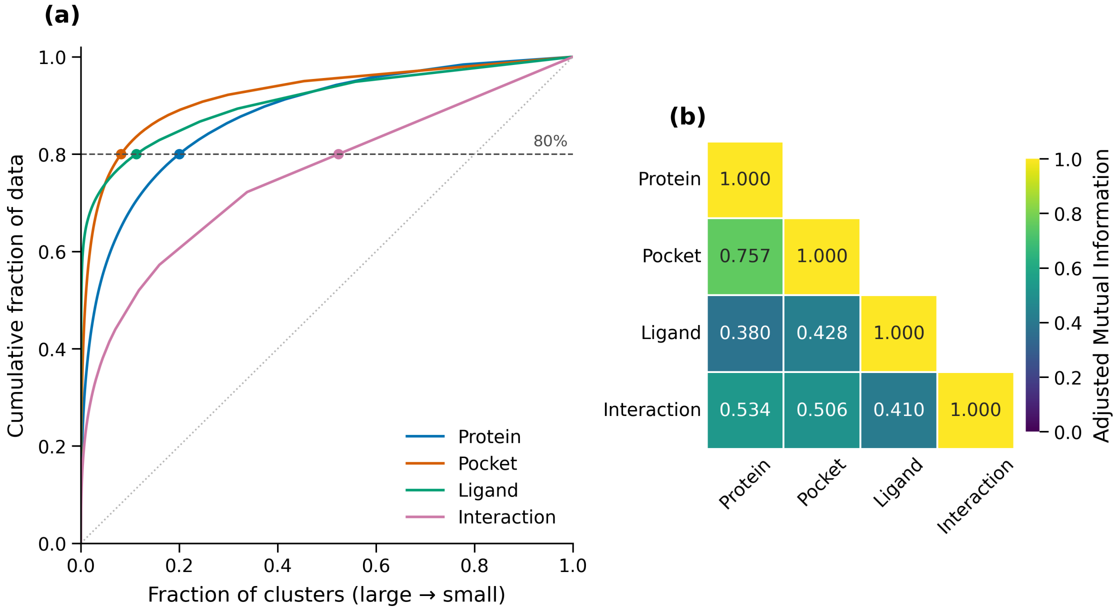


Figure S3: **Hierarchical clustering of CROWN entries.** (a) Cumulative data coverage for each clustering. Clusters are ordered from largest to smallest, and the cumulative fraction of data points is plotted against the fraction of clusters included. Filled circles mark the point at which each curve reaches 80% coverage; the dotted diagonal indicates the line of equal cluster sizes. Curves bowing further toward the upper left reflect more top-heavy clusterings, in which a small fraction of clusters accounts for most of the data. (b) Pairwise agreement between clustering schemes, reported as the Adjusted Mutual Information computed on the 70% similarity cluster labels.
